# Hi-C2B: Optimised detection of chromosomal contacts within synchronised meiotic *S. cerevisiae* cells

**DOI:** 10.1101/2023.11.21.565821

**Authors:** Ellie M. Wright, S. Schalbetter, Matthew J. Neale

## Abstract

Hi-C, a genome-wide chromosome conformation capture assay is a powerful tool used to study three-dimensional genome organisation by converting physical pairwise interactions into counts of pairwise interaction. To study the many temporally regulated facets of meiotic recombination in *S. cerevisiae* the Hi-C assay must be robust such that fine- and wide-scale comparisons between genetic datasets can be made. Here we describe an updated protocol for Hi-C (Hi-C2B) that generates reproducible libraries of interaction data with low noise and for a relatively low cost.

## 1. Introduction

Chromosome conformation capture methodologies based on “3C”—a “one–to–one” approach for studying contact frequencies between known genomic sequences—have advanced our understanding of chromosome and nuclear organisation in many organisms *(1–10, 14)*. Prior to the evolution of 3C, chromosome structure was studied via a combination of electron microscopy, immunofluorescence microscopy and the global, spatial patterning of proteins of interest determined by chromatin immunoprecipitation (ChIP). Now, chromosome conformation can also be analysed with respect to the underlying genomic sequence unveiling a wealth of information about the structural properties and spatial conformations of chromosomes within individual or populations of cells.

Early discoveries attributed to the 3C assay include identification of “flexible” chromosomes within *S. cerevisiae (1)*; observations of enhancer–promoter interactions for gene regulation at the β-globin locus *(2)*, furthered by studies that showed enhancer–promoter interactions require tissue-specific transcription factors *(3–4)*. Architectural chromatin loops have been uncovered as stable chromosome structures mediated, in mammals, by a partnership between ubiquitously expressed CTCF protein and the cohesin complex *(5–9)*. On the back of such discoveries, and a greater accessibility and availability of high-throughput DNA sequencing technologies, development of many 3C-related techniques—including Hi-C—have transpired. Each 3C-based method customises the technique to provide independent, detailed documentation of the spatial organisation of chromosomes *(10)*.

To capture chromosome conformation via 3C-related methodologies, the same initial approach is employed. Briefly, intact cells are isolated and subjected to treatment with formaldehyde in order to cross link physical DNA–protein and protein–protein interactions occurring in three-dimensional space *(1)*. Cross-linked material is digested with a 6- or 4-bp cutting restriction enzyme, the latter increasing the spatial resolution of resulting data. Subsequent adaptations of the Hi-C method have also employed the use of MNase (Micro-C) to generate Hi-C interaction maps at nucleosome resolution *(11)*. In all methods, fragmented DNA is subjected to in situ ligation prior to cross link reversal. At this stage, a 3C “template” is obtained which contains both linear and circular polymeric concatemers made from genomic segments reshuffled with respect to their spatial proximity at the time of fixation *(10)*. Following generation of a 3C “template”, 3C-based techniques differ in their detection and quantification of 3C ligation junctions *(10)*.

Like classic 3C, Hi-C employs one of a number of restriction enzymes to digest cross-linked DNA to generate 5’ overhangs that are subsequently filled with a biotin-labelled nucleotide (Fig. 1) *(12)*. Blunt-end fragments are then ligated to produce products where the ligation junction is marked with the biotin label, referred to as the 3C template *(12)*. A Hi-C library is then created by mechanical shearing followed by enrichment steps before paired-end high-throughput sequencing, and mapping reads to a reference genome *(12)*. The result: a two-dimensional matrix of the sequence-specific interactions occurring within the three-dimensional nuclear space *(12)*.

**Fig 1.**
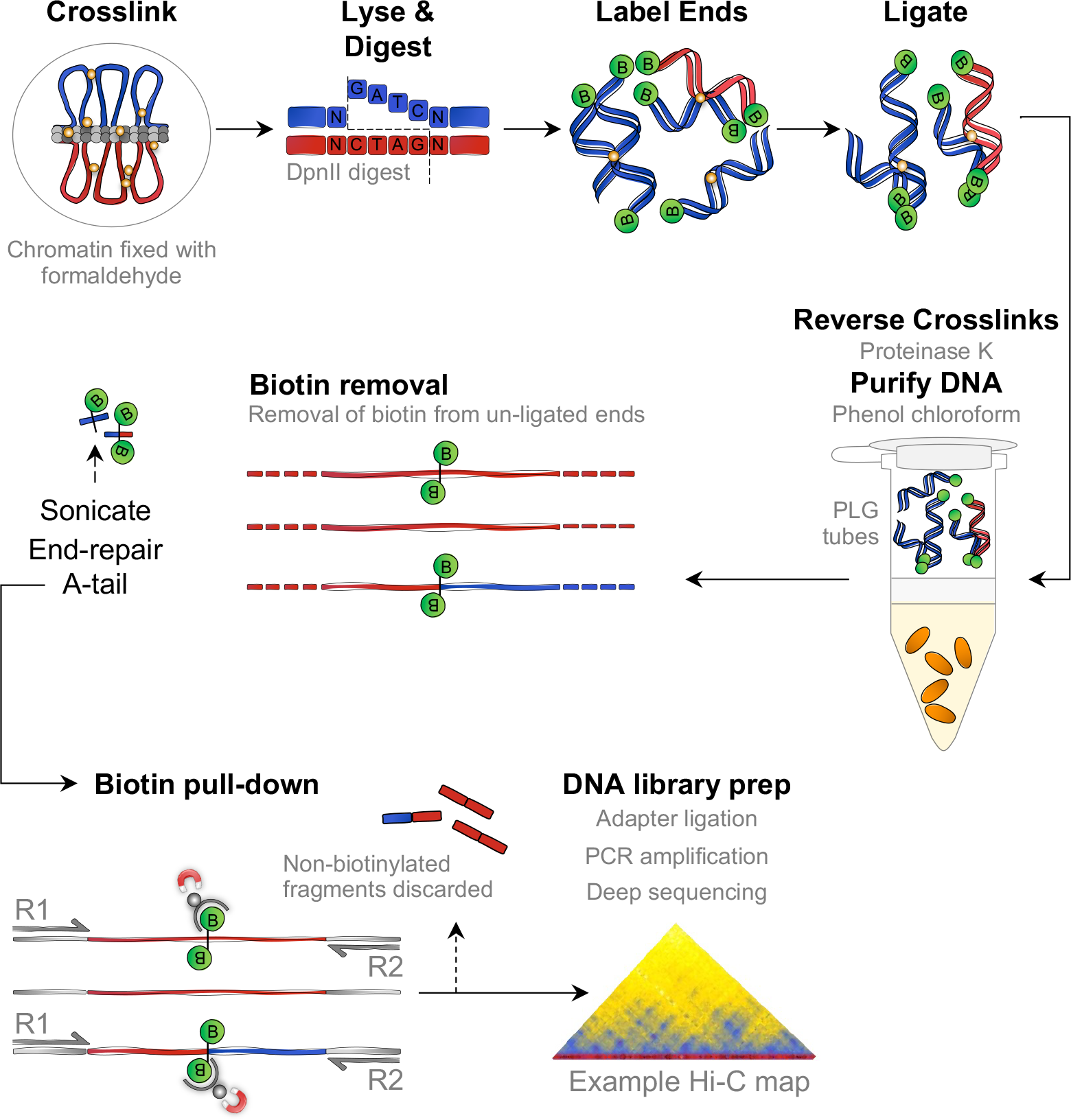
Schematic outlining the various steps of Hi-C library preparation. Chromatin is fixed with formaldehyde to capture DNA–DNA and DNA–protein interactions. Cells are lysed and chromatin solubilised before the DNA is digested with DpnII to generate 5’-GATC overhangs. Overhangs are filled with nucleotides including biotin-14-dCTP (green filled circles). Blunt ends are ligated together and cross-links are reversed via proteinase K treatment. DNA is extracted with phenol:chloroform. Biotin is removed from unligated ends, chimeric DNA fragments are sheared, sonicated ends are repaired and subject to to poly(A)-tailing. Ligation junctions are enriched via streptavidin-coated beads which have a high affinity for the incorporated biotin. Non-biotinylated fragments are discarded before adapter ligation, PCR amplification, deep sequencing and mapping (see Methods for full details).

**Fig 2.**
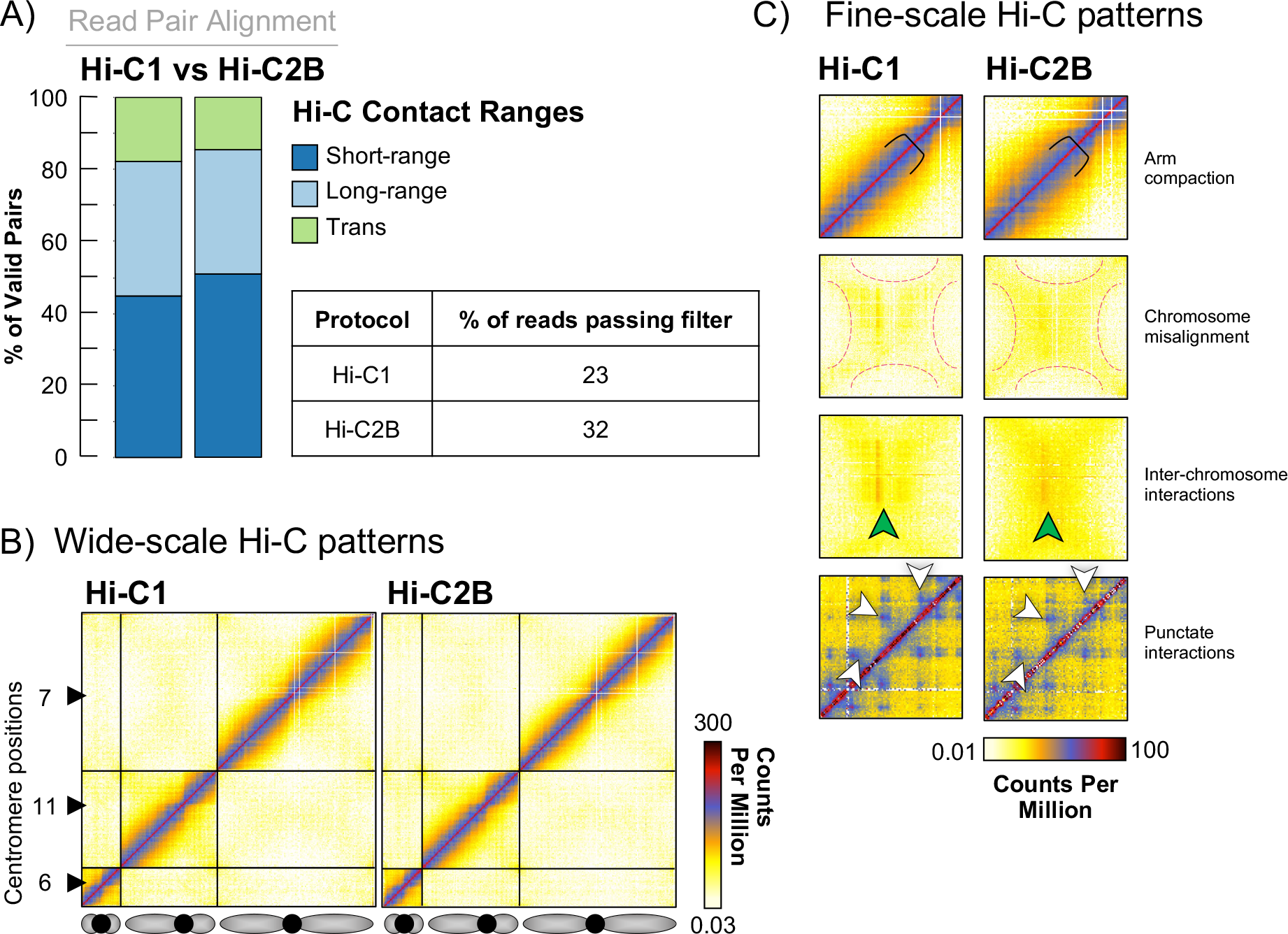
Comparisons between Hi-C1 and Hi-C2B data output. **A)** Fragment size distribution of short- and long-range intra-chromosomal (cis) contacts versus long-range inter-chromosomal (trans) contacts as reported by the Hi-C Pro pipeline *(14)* for libraries assayed by Hi-C1 or Hi-C2B methodologies. The table shows the percentage of reads that pass all Hi-C Pro filters as a fraction of the starting read value (all pairs). **B)** Hi-C interaction maps from meiotic *S. cerevisiae* cells prepared by Hi-C1 or Hi-C2B. Assessment of wide-scale patterns of Hi-C interaction highlight the similarity in output data from either methodology. Hi-C contact maps of chromosome 6, 11, and 7 are plotted at 5 Kb resolution. The black arrows indicate positions of the centromeres. **C)** Assessment of known chromosome conformations demonstrates similarities in the fine-scale Hi-C interaction patterns from libraries prepared by Hi-C1 or Hi-C2B. Arm compaction: axial compaction is indicated by the width of the main diagonal relative to the fixed-width black clamp. Chromosome misalignment: dashed pinked lines outline the inter-chromosomal cross-shape structures that manifest due to chromosome misalignment (present in meiotic recombination mutants for example). Inter-chromosome interactions: enhanced domains of inter-chromosomal contacts indicated by green arrowheads. Punctate interactions: focal grid-like patterns that arise from loops generated by meiotic cohesin (Rec8) *(18)*; prominent examples are indicated by white arrowheads. Hi-C contact maps from meiotic *S. cerevisiae* cells are plotted at 2 Kb resolution.

Hi-C, the genome-wide adaptation of 3C, was introduced more than a decade ago (Hi-C1) *(1)*. Hi-C2, described as an optimised method for the production of high-resolution chromosome conformation, includes a number of changes to the Hi-C1 protocol, particularly to the blunt-end ligation reaction *(13)*: 1) removal of additional chromatin solubilisation by SDS prior to blunt-end ligation, with the aim to preserve nuclear structure thus reducing the occurrence of random ligation events; 2) reduced reaction length; 3) abolition of large-scale dilution during ligation which was included in the original Hi-C protocols to avoid random intermolecular ligation events. Hi-C1 *in situ* ligation reactions are diluted 15-fold whereas the Hi-C2 protocol documents just a 2-fold dilution; finally, 4) Hi-C2 includes a molecular crowding agent, polyethylene glycol (PEG) during ligation.

The methodological adaptations of Hi-C2 yielded a protocol with smaller processing and reaction volumes—thereby, dramatically reducing the overall cost of Hi-C library preparation *(13)*. In addition, smaller processing volumes make handling of samples much easier such that more samples can be processed in parallel. Performing Hi-C1 restricted handling to a maximum of two samples per assay. However, approximately eight samples can be processed in a single run of Hi-C2. Despite these benefits, we found that employing a Hi-C2-like protocol in *S. cerevisiae* generally yielded libraries with high levels of inter-chromosomal (trans) Hi-C interactions, indicative of random ligation.

Here we present a composite chromosome conformation capture assay that combines beneficial aspects of both the Hi-C1 and Hi-C2 protocols in order to generate high quality, reproducible *S. cerevisiae* Hi-C interaction maps with high yield, throughput and relatively low cost. For a summary of changes see **Table 1**.

**Table 1.** Summary table of changes between Hi-C1 and Hi-C2B protocols.

| Protocol | Hi-C1 | Hi-C2B |
| --- | --- | --- |
| Spheroplast & lyse | 35°C, 15 min, invert every 5 min;<br>0.1% SDS, 65°C, 10 min | 35°C, 10 min, invert every 2–3 min;<br>0.01% SDS, 65°C, 5 min |
| Digest Enzyme | 2.07 U/μL DpnII | 0.92 U/μL DpnII |
| Label | 37°C, 2 hours | 23°C, 4 hours |
| Solubilise | 1.5% SDS, 65°C, 20 min | NA |
| Blunt end ligation | 15-fold dilution;<br>10X NEB T4 DNA ligase buffer (w/o PEG), 16°C, 8 hours;<br>Split across 6x 2 mL tubes | 2-fold dilution;<br>10X NEB T4 DNA ligase buffer (w/o PEG), 16°C, 4 hours;<br>Split across 16x PCR tubes |
| Extract DNA | 2-step DNA extraction;<br>Precipitate whole ligation reaction in EtOH, 4°C, overnight (large volume);<br>Spin down & dissolve the pellet in TE. Proteinase K crosslink reversal, 65°C, 2 hours;<br>PCI DNA extraction and precipitation, 4°C, 2 hours (small volume) | PCI DNA extraction and precipitation of whole ligation reaction, 4°C, overnight |
| Remove biotin | dATP, dTTP, dCTP, dGTP used in reaction | dATP, dGTP <u>only</u> used in reaction |

## 2. Materials

Media solutions are prepared using double distilled water, dH_2_O, and autoclaved. Prepare and store all reagents at room temperature (unless indicated otherwise). Materials are listed in the first section in which they are used. Product codes are denoted by # followed by a number.

### 2.1. Meiotic Cell Cultures

1. Amino acid supplement (AAHLTU) 200X: Weigh 2 g of the following powders Adenine hemisulfate salt (#A3159), L-Arginine monohydrochloride (#A5131), L-Histidine monohydrochloride monohydrate (#H8125), L-Leucine (#L8000), L-Tryptophan (#T0245), Uracil (#U0750).
2. YPD (2%) media: 1% (w/v) Yeast extract, 2% (w/v) Peptone, 2% (v/v) Glucose, 20%, 0.0137% (w/v) Adenine + Uracil (AU) supplement.
3. BYTA media: 1% (w/v) Yeast extract, 2% (w/v) Tryptone, 1% (w/v) Potassium Acetate, 50 mM Potassium Phthalate, 0.001% (v/v) Antifoam 204 (Sigma #A8311).
4. Sporulation media (SPM): 0.3% (w/v) Potassium acetate, 0.02% (w/v) Raffinose, 0.004% (v/v) AAHLTU, 200X, 0.001% (v/v) Antifoam solution.
5. Copper (II) sulphate (250mM): Weigh 0.39 g CuSO_4_ powder (Sigma #C1297) and dissolve in 10 mL dH_2_O.

### 2.2. Fixation

1. 37% Formaldehyde (FA): Formaldehyde solution (Sigma #F8775).
2. 2.5 M Glycine: Weigh 75 g glycine powder (Sigma #G5417) and dissolve in 1L dH_2_O. Filter to sterilise.
3. Dry ice.

### 2.3. Spheroplast, Lyse, Digest

1. Spheroplast buffer: 50 mM Tris-HCl (pH 7.5), 1 M Sorbitol. Store at 4°C.
2. 2-Mercaptoethanol (Sigma Aldrich #M6250).
3. 100T Zymolyase stock (50 mg/mL): Weigh 0.1 g Zymolyase 100T powder (#120493-1) and dissolve in 1.4 mL 50% glycerol, 0.1 mL 1 M NaHPO_4_ buffer (pH 7.2), 0.5 mL 2 M sucrose. Vortex to dissolve, store at -20°C. Dilute to 5 mg/mL by taking 200 μL of the 50 mg/mL stock and diluting in 800 μL 50% glycerol. Store at -20°C.
4. NEBuffer™ 1X r3.1: Dilute 10X NEBuffer™ r3.1 (#B6003S) in dH_2_O. Keep on ice.
5. 1% SDS: Dilute 10% SDS in dH_2_O.
6. 10% Triton X-100: Dilute Triton X-100 in dH_2_O.
7. NEB™ DpnII (#R0543L). Store at -20°C.

### 2.4. Fill-in, Blunt-end Ligate, Reverse Crosslinks

1. NEBuffer™ 10X r3.1 (#B6003S). Store at -20°C.
2. Thermo Scientific™ 10 mM dATP, dTTP, dGTP: Individually dilute 10 μL of 100 mM dNTPs (#10520651) with 90 μL of Tris-EDTA (TE). Store at -20°C.
3. Invitrogen™ Biotin-14-dCTP: 0.4 mM biotin-14-dCTP (#10022582), 100 mM Tris-HCl (pH 7.5), 0.1 mM EDTA.
4. NEB™ Klenow DNA Polymerase I (5U/μL) (M0210L). Store at -20°C.
5. Eppendorf™ ThermoMixer™ (#5382000031).
6. NEB™ T4 DNA Ligase Buffer (10X): 50 mM Tris-HCl (pH 7.5), 10 mM MgCl2, 1 mM ATP, 10 mM DTT (#B0202S).
7. Invitrogen™ T4 DNA Ligase Buffer (5X): 250 mM Tris-HCl (pH 7.6), 50 mM MgCl2, 5 mM ATP, 5 mM DTT, 25% (w/v) polyethylene glycol-8000 (included with: #10443242).
8. Bovine Serum Albumin (BSA) 10 mg/mL: Weigh 10 mg BSA (#B9001S) and dissolve in 1 mL dH_2_O to make 10 mg/mL.
9. Invitrogen™ T4 DNA Ligase (5U/μL) (#10443242). Store at -20°C.
10. Eppendorf™ 2 mL safe-lock tubes (#10430423).
11. Proteinase K (10 mg/mL): Weigh 0.1 g of proteinase K powder (Fisher Scientific #10407583) and dissolve in 10 mL of 20 mM Tris-HCl 50% glycerol solution. Store at -20°C.

### 2.5. Extract, Precipitate and Quantify DNA

1. Phenol – chloroform – isoamyl alcohol mixture (#77617). Store at 4°C.
2. 5PRIME Phase Lock Gel (PLG) tubes – light (#2302820).
3. 3 M sodium acetate, pH 5.2: Weigh 246.1 g of sodium acetate and dissolve in 800 mL of dH_2_O. Adjust the pH to 5.2 with glacial acetic acid. Allow the solution to cool to RT before use.
4. Millipore Amicon® Ultra 0.5 mL centrifugal filters (#UFC500396).
5. Tris Low-EDTA (TLE): 10 mM Tris-HCl (pH 8.0), 0.1 mM EDTA.
6. Ribonuclease A (RNase A) 10mg/mL: Weigh 0.1 g of RNase A powder (Sigma Aldrich #R4875) and dissolve in 9 mL of 10 mM sodium acetate buffer (pH 5.2). Heat to 100°C for 15 minutes and allow to cool to room temp before use. Adjust to pH 7.4 by adding 0.1 mL of 1M Tris-HCl (pH 7.4). Store at -20 °C.
7. Qubit™ Fluorometer (purchased from Thermo Scientific™).
8. Invitrogen™ Qubit™ dsDNA high sensitivity assay kit (#Q32851).

### 2.6. Remove Dangling-ends

1. NEBuffer™ 10X r2.1 (#B6002S). Store at -20°C.
2. 10 mM dATP, dGTP: Prepared as described in 2.4.
3. NEB™ T4 DNA Polymerase I (3U/μL) (#M0203L). Store at -20°C.

### 2.7. Fragment, Repair Broken-ends

1. Covaris microTUBE: AFA^®^ fibre pre-slit snap-cap (130 μL) (#520045).
2. Covaris instrument: We use a low-throughput focused-ultrasonicator, M220 (#500295).
3. 100 mM dNTP mix: In a single Eppendorf, mix 100 μL of each dNTP (#10520651). Store at -20°C.
4. NEB™ T4 Polynucleotide Kinase (PNK): Store at -20°C (#M0201S).

### 2.8. Enrich, Ligate, PCR Amplify

1. DNA LoBind Tube 1.5 mL (#022431021).
2. Invitrogen™ Dynabeads^®^ MyOne™ Streptavidin C1 beads (#65001).
3. Tween Wash Buffer (TWB): 50 mM Tris-HCl (pH 8.0), 0.5 mM EDTA, 1 M NaCl, 0.05% (v/v) Tween.
4. Invitrogen™ Dynal MPC™: Magnetic particle concentrator.
5. Binding Buffer (2X): 10 mM Tris-HCl (pH 8.0), 1 mM EDTA, 2 M NaCl.
6. Binding Buffer (1X): Dilute 5 mL 2X BB in 5 mL dH_2_O. Make fresh for each experiment and store on ice.
7. Klenow Fragment (3’–5’ exo-) (#M0212S). Store at -20°C.
8. T4 DNA Ligase Buffer (1X): Dilute Invitrogen™ T4 DNA Ligase Buffer (5X) in dH_2_O. Make fresh for each experiment and store on ice.
9. NEXTFLEX^®^ DNA Barcodes (#NOVA-514103).

### 2.9. Clean, Quantify, Sequence, Map

1. Beckman Coulter™ Agencourt AMPure^®^ XP (#A63881). Store at 4°C.
2. Bioanalyser instrument: We use Agilent 2100 Bioanalyzer Instrument (#G2939BA).
3. Agilent High Sensitivity DNA Kit (#5067-4626).
4. Illumina NexSeq 500/550 High Output Kit v2.5 (75 cycles) (#20024906).
5. Linux workstation running Hi-C Pro *(15)* (version 3.1.0).

## 3. Methods

The protocol presented here is optimised for the detection of intra- and inter-chromosomal contacts from highly synchronous *S. cerevisiae* cells. Both cycling and sporulating budding yeast cells can be investigated using the Hi-C2B assay. We combine methodological steps from Hi-C1 and Hi-C2 in order to improve the recovery and resolution of *S. cerevisiae* Hi-C libraries. A schematic outlining the various steps of the Hi-C library preparation is shown in **Fig. 1**.

### 3.1. Meiotic Cell Cultures

1. From a –80°C glycerol stock, streak out yeast cells onto a YPD plate. Incubate 2 days at 30°C.
2. In a glass culture tube, inoculate 4 mL YPD liquid with a single colony taken from the YPD plate. Incubate 24 hours at 30°C with shaking (250 rpm), until the culture is saturated (∼OD 20).
3. In a conical flask, inoculate the saturated culture in BYTA pre-sporulation media to a final OD of 0.3. Incubate for 16–18 h at 30°C with shaking (250 rpm) (*see* Note 1).
4. Collect cells by centrifugation (4 minutes, 4,000 *g*, room temp) and drain supernatant. Wash with water by sealing the tube and shaking vigorously (*see* Note 2). Pellet cells again, drain, and resuspend with SPM. Return to a conical flask and incubate at 30°C with shaking (250 rpm) for the duration of the time course (*see* Note 3).

### 3.2. Fixation

1. Transfer 8 mL (20–30 OD_600_ units) of synchronously sporulating *S. cerevisiae* cells to a 50 mL Falcon tube.
2. Add 648 μL 37% formaldehyde (∼3% final concentration). Incubate for 20 minutes at 30°C with shaking (250 rpm).
3. Quench formaldehyde by adding 1.3 mL of 2.5 M glycine (∼0.35 M final) and incubate for a further 5 minutes at 30°C with shaking (250 rpm) (*see* Note 4).
4. Collect cells by centrifugation (2 minutes, 2,500 *g*, 4°C) and aspirate supernatant. Wash with 1.4 mL water by resuspending material using a pipette, do this slowly to avoid making bubbles.
5. Split the suspension into four separate 1.7 mL Eppendorf tubes (∼350 μL per tube) (*see* Note 5).
6. Collect cells, again by centrifugation using a table top centrifuge (1 minute, 14,000 *g*, room temp). Carefully remove the supernatant using a pipette and snap freeze the pellets on dry ice (*see* Note 6).

### 3.3. Spheroplast, Lyse, Digest

1. Thaw cell pellets on ice for 1–2 hours.
2. Wash cells in 1 mL ice-cold spheroplasting buffer by pipetting up and down (do not vortex, the cells are fragile at this stage).
3. Collect cells by centrifugation using a table top centrifuge (2 minutes, 2,500 *g*, room temp). Carefully remove supernatant using a pipette (*see* Note 7).
4. Resuspend cells in 1 mL spheroplasting buffer. Add 2 μL 50 mg/mL zymolyase 100T (100 μg/mL final) and 10 μL 2-mercaptoethanol and mix thoroughly.
5. Incubate for 10 minutes at 35°C, inverting slowly after five minutes.
6. Collect cells by centrifugation (2 minutes, 2,500 *g*, room temp) then resuspend the pellet in 500 μL ice-cold 1X NEB 3.1 restriction enzyme buffer (*see* Note 8).
7. Collect cells as described in 6. Resuspend the pellet in 360 μL ice-cold 1X NEB 3.1 restriction enzyme buffer.
8. Solubilise chromatin by adding 3.8 μL 1% SDS and mixing gently by pipetting up and down to avoid making bubbles. Incubate at 65°C for 5 minutes.
9. Cool mixture by placing it on ice for 5 minutes immediately after incubation with SDS.
10. When the mixture has cooled, add 43 μL of 10% Triton X-100 and invert gently, 2–3 times to quench SDS. Incubate for 15 minutes at 37°C, 300 rpm (*see* Note 9).
11. Add 8 μL DpnII, mix thoroughly by inversion and incubate overnight in a thermomixer at 37°C, 300 rpm. The final volume at this stage will be ∼460 μL.

### 3.4. Fill-in, Blunt-end Ligate, Reverse Crosslinks

1. Incubate at 65°C for 20 minutes in order to deactivate DpnII.
2. Fill-in DNA ends by adding 2 μL water, 6 μL 10X NEB 3.1 restriction enzyme buffer, 1.5 μL 10 mM dATP, dGTP and dTTP, 37.5 μL 0.4 mM biotin-14-dCTP, and 5 U/μL Klenow fragment DNA polymerase I. Mix by carefully pipetting and gentle inversion (3–4 times).
3. Incubate mixture at 23°C for 4 hours with interval agitation in a ThermoMixer (*see* Note 10).
4. For blunt-end ligation, make a reaction mixture containing 120 μL 10% Triton X-100, 120 μL 10X ligation buffer (NEB), 12 μL 10 mg/mL BSA, 363 μL dH_2_O, 50 μL 5U/μL T4 DNA Ligase (Invitrogen). Total will be 665 μL (*see* Note 11).
5. Dilute the sample 2-fold by adding the ligation reaction mixture to the fill-in reaction tube and mixing fully by inverting the tube 3–4 times.
6. Split volume across sixteen 0.2 mL PCR tubes (∼70 μL per tube) (*see* Note 12).
7. Incubate at 16°C for 4 hours inverting slowly after thirty minutes.
8. Pool the ligation reaction into a single 2 mL safe-lock tube (pulse spin to recover material that may be in the lid).
9. Add 100 μL of 10 mg/mL proteinase K and incubate at 65°C overnight to reverse crosslinks.

### 3.5. Extract, Precipitate and Quantify DNA

1. Cool mixture by placing it on ice for ∼20 minutes.
2. Transfer half of the mixture (∼600 μL) to a second 2 mL safe-lock tube (two tubes per sample).
3. Add an equal volume of PCI (25:24:1) to each tube and vortex heavily. Transfer the white, milky solution to a pre-spun 2 mL phase-lock tube (2 per sample).
4. Centrifuge at room temp for 5 minutes, 14,000 *g*.
5. Transfer the aqueous phase containing DNA, to pre-chilled 1.7 mL Eppendorf tubes (2 per sample) containing 50 μL 3M NaOAc and 1 mL 100% EtOH (pre-chilled).
6. Store on dry ice for a few minutes or at -20°C for at least 2 hours to precipitate DNA (*see* Note 13).
7. Collect the DNA precipitate by centrifugation at 4°C for 30 minutes, 16,000 *g* and aspirate supernatant (pulse spin to remove residual EtOH).
8. Wash the pellet with 1 mL ice-cold 70% EtOH.
9. Repeat step 7 (*see* Note 14).
10. Air dry the pellet for ∼20 minutes.
11. Dissolve the pellet in 250 μL 1X TLE (pH = 8.0) for ∼30 minutes.
12. Pool into a single Amicon 30 kDa column and centrifuge at room temp, 14,000 *g* until volume reaches ∼100 μL (*see* Note 15).
13. Wash with 400 μL 1X TLE (pH = 8.0) and repeat spin until volume reaches ∼100 μL.
14. Flip the Amicon 30 kDa column containing the concentrated sample into a new collection tube, spin at room temp for 2 minutes, 14,000 *g*.
15. Transfer sample to a 1.7 mL Eppendorf tube and determine the volume (usually ∼100–130 μL).
16. Add 1:100 μL of 1 mg/mL RNAse A (typically ∼1–1.3 μL) and incubate at 37°C for 30 minutes.
17. Measure the DNA concentration of the Hi-C sample using the Qubit dsDNA HS Assay Kit and proceed to the removal of “dangling ends” (*see* Note 16).

### 3.6. Remove Dangling-ends

1. Remove biotin-14-dCTP from un-ligated ends by adding 12 μL 10X NEB 2.1 restriction enzyme buffer, 0.3 μL 10 mM dATP, 0.3 μL 10 mM dGTP, 12 μL 300 U/μL T4 DNA Polymerase (*see* Note 17).
2. Incubate the mixture at 20°C for 4 hours and inactivate the enzyme by incubating at 75°C for 20 minutes.
3. Transfer the mixture into a single Amincon 30 kDa column and centrifuge at room temp, 14,000 *g* until volume reaches ∼100 μL.
4. Wash column with 400 μL 1X TLE (pH = 8.0) and repeat spin until volume reaches ∼130 μL.
5. Flip the Amicon 30 kDa column containing the concentrated sample into a new collection tube, spin at room temp for 2 minutes, 14,000 *g*. Determine the volume in the column.
6. Bring the Hi-C sample to 130 μL with 1X TLE and load into a Covaris microTUBE.

### 3.7. Fragment, Repair Broken-ends

1. Load the microTUBE containing the Hi-C sample into a Covaris M220.
2. Run the following program (*see* Note 18):
  1. Duty factor: 20%
  2. Peak power: 50W
  3. Cycles per burst: 200
  4. Temperature: 20°C
  5. Run time: 240 seconds
3. Transfer the sonicated Hi-C sample to a 0.2 mL PCR tube.
4. To repair broken DNA ends, add to the sample 15 μL 10X ligation buffer (NEB), 1.5 μL 100 mM dNTP mix (25 mM each), 5.4 μL 3U/μL T4 DNA Polymerase, 5.4 μL 10U/μL T4 Polynucleotide Kinase, 1.1 μL 5U/μL Klenow DNA Polymerase I. Make the volume up to 30 μL with water and mix.
5. Incubate at 20°C for 30 minutes and inactivate enzyme activity by incubating at 75°C for 20 minutes. Maintain at 4°C.

### 3.8. Enrich, Ligate, PCR Amplify

1. Vortex Streptavidin C1 beads and transfer 10 μL into a 1.7 mL DNA LoBind tube (*see* Note 19).
2. Wash beads with 100 μL of Tween Wash Buffer (TWB) by pipetting up and down.
3. Incubate the beads with the buffer at room temp for 3 minutes with rotation.
4. Collect the beads against a magnetic particle concentrator at room temp for 1 minute and remove and discard the supernatant.
5. Resuspend the beads in 100 μL of TWB and transfer to a new LoBind tube.
6. Repeat step 4.
7. Resuspend the beads in 150 μL of 2X Binding Buffer (BB) and add to the end repair reaction. Pipette up and down to mix, avoid making bubbles.
8. Incubate at room temp for 1 hour with rotation.
9. Repeat step 4.
10. Resuspend beads in 400 μL 1X BB and transfer to a new LoBind tube. Gently invert to mix.
11. Repeat step 4.
12. Resuspend beads with 100 μL 1X TLE (pH 8.0) and transfer to a new LoBind tube.
13. Repeat step 4.
14. Resuspend beads in 41 μL 1X TLE (pH 8.0).
15. Digest unligated biotinylated dangling ends by adding 5 μL 10X NEB 2.1 restriction enzyme buffer, 1 μL 10 mM dATP, 3 μL Klenow DNA Polymerase (3’–5’ exo-).
16. Incubate at 37°C for 30 minutes and inactivate the enzyme activity by incubating at 65°C for 20 minutes. Cool mixture by incubating at 4°C.
17. Transfer the reactions to a new LoBind tube, then repeat step 4.
18. Resuspend beads in 400 μL 1X T4 DNA ligation buffer.
19. Repeat step 4.
20. Resuspend beads in 40 μL 1X T4 DNA ligation buffer.
21. Ligate a unique NEXTFLEX^®^ barcoded adaptor by adding 2 μL 5X T4 DNA ligation buffer (Invitrogen), 3 μL 25 μM NextFlex barcoded adaptor, 3 μL 5X T4 DNA ligase (Invitrogen), 2 μL water and mix.
22. Incubate at 22°C for 2 hours (*see* Note 20).
23. Repeat step 4.
24. Was beads twice with 400 μL TWB by repetition of step 4 in between washes. Incubate at room temp for 5 minutes with rotation.
25. Repeat step 4.
26. Resuspend beads in 200 μL 1X BB and transfer to a new LoBind tube.
27. Repeat step 4.
28. Resuspend beads in 200 μL 1X TE and transfer to a new LoBind tube.
29. Repeat step 4.
30. Resuspend beads in 10 μL 1X TE and transfer to a new LoBind tube.
31. Minimally amplify the Hi-C library by adding 40 μL 2X NEXTFLEX^®^ DNA PCR Master Mix, 6.7 μL NEXTFLEX^®^ Primer Pair Mix, and 110 μL water to the DNA on beads and mix.
32. Divide the reaction between 4 PCR tubes (∼41 μL per tube) and run the following programme:
  a. 98°C, 2 minutes
  b. 12–14 cycles of:
    1. 98°C, 30 seconds
    2. 65°C, 30 seconds
    3. 72°C, 1 minute
  c. 72°C, 4 minutes
33. Collect the beads against a magnetic particle concentrator at room temp for 1 minute and **retain** the supernatant by pooling into a single 1.7 mL Eppendorf tube. The total volume will be ∼150 μl. **This is the Hi-C sample**. Store at -20°C before proceeding to the next step.

### 3.9. Clean, Quantify, Sequence, Map

To remove primer dimers, purify the amplified Hi-C library from the supernatant using Ampure XP beads. Allow the Ampure XP mixture to come to room temp (for ∼30 minutes) and vortex heavily prior to use. Pre-warm 130 μL (per sample) 1X TLE buffer to 65°C.

1. Add 1.1x Ampure XP beads (∼165 μL) to the Hi-C sample.
2. Vortex and lightly spin to remove mixture from the lid.
3. Incubate at room temp for 10 minutes.
4. Collect the beads against a magnetic particle concentrator at room temp for 5 minutes and remove and discard supernatant.
5. Wash beads twice by adding 1 mL 70% EtOH to the tube, incubate at room temp for 1 minute, then aspirate. Repeat (*see* Note 21).
6. Elute DNA by resuspending the DNA/beads in 100 μL 1X TLE preheated to 65°C (see above).
7. Incubate at room temp for 10 minutes.
8. Add 1.1x Ampure XP beads (110 μL) and repeat steps 2–5 with the eluent/bead mixture.
9. Repeat step 6, but resuspend in 30 μL 1X TLE preheated to 65°C.
10. Incubate at room temp for 10 minutes.
11. Repeat step 4, however, this time **<u>keep</u>** the supernatant (Hi-C library) and transfer to a new 1.7 mL Eppendorf tube labelled with the library name/number.
12. Assess the size and concentration of DNA using the Agilent 2100 Bioanalyzer system.
13. Sequence Hi-C libraries using paired-end reads of at least 42 bp (using, for example, a NextSeq sequencing kit) (*see* Note 22).
14. Sequenced Hi-C data can be mapped using Hi-C Pro using standard parameters (https://github.com/nservant/HiC-Pro) (*see* Note 23).

## 4. Notes

1. Under the control of the *CUP1* promoter, transcriptional induction of *IME1* depends solely on the presence of copper ions *(15)*. As such, whereas acetate-based pre-sporulation media has been shown to boost sporulation compared to glucose-based in non-inducible cells, acetate-based pre-growth is dispensable for sporulation induced from the *CUP1* promoter *(15–17)*. An alternative pre-sporulation medium that is compatible with the Hi-C2B assay is low glucose YP (as described in *15*). Cells are prepared as described in 3.1.1 with a few additional steps. Following step 2, the saturated culture is transferred to fresh YPD media in either a glass culture tube or conical flask (depending on the final volume) to a final OD of 0.1 and grown to exponential phase for a few hours. In place of step 3, the exponential culture should be inoculated in reduced glucose YPD pre-sporulation media (we use 1%) to a final OD of 0.05. As above, incubate for 16–18 h at 30°C with shaking (250 rpm), until the culture reaches OD 10–12. Proceed to step 4. For a more detailed protocol please ref 15.
2. We find that heavily vortexing the cellular material for a full resuspension at this stage improves synchronous release into meiosis. Water should be stored at room temp.
3. For strains where *IME1* is placed under the control of the *CUP1* promoter, after 2 hours in sporulation medium, copper (II) sulphate (50 mM) is added to induce *IME1* expression and initiate sporulation synchronously.
4. Holding the mixture on ice for ∼10 minutes helps to further quench fixed cells resulting, in our experience, in an improved library yield.
5. Hi-C2B features a 2-fold reduction in starting material. While the total volume remains the same, the concentration of reagents in proportion to the cellular material is higher in Hi-C2B. Reducing the amount of starting material appears to improve the resolution of Hi-C libraries.
6. Snap frozen, fixed cell pellets should be stored at -80°C.
7. Cell pellets are relatively unstable at this point so care should be taken when removing the supernatant. The same applies for step 6.
8. Dilute 10X NEB 3.1 restriction enzyme buffer in Milli-Q water and chill on ice.
9. Short incubation of the solubilised material with Triton X-100 prior to the addition of the enzyme improves DpnII efficiency—inferred by improved resolution of final Hi-C libraries—likely due to a reduction in SDS-associated inhibition of enzyme activity.
10. Eppendorf ThermoMixer Settings: 23°C, 4 hours; 900 rpm for 10 seconds every 5 minutes.
11. Despite using T4 DNA Ligase from Invitrogen, it is important to substitute the 5X Ligation Buffer for the 10X NEB variety which does not contain polyethylene glycol (PEG). While previous methods have reported the use of ligase buffer containing PEG for generation of Hi-C libraries *(16)*, replacement of PEG-containing (Invitrogen) with non-PEG-containing (NEB) ligation buffer minimises non-specific intermolecular ligation efficiency, thereby favouring intramolecular ligation. This alteration is crucial for the generation of high-resolution Hi-C contact maps that are not swamped with artefactual intermolecular interactions (see Fig. 3 for further details).

**Fig 3.**
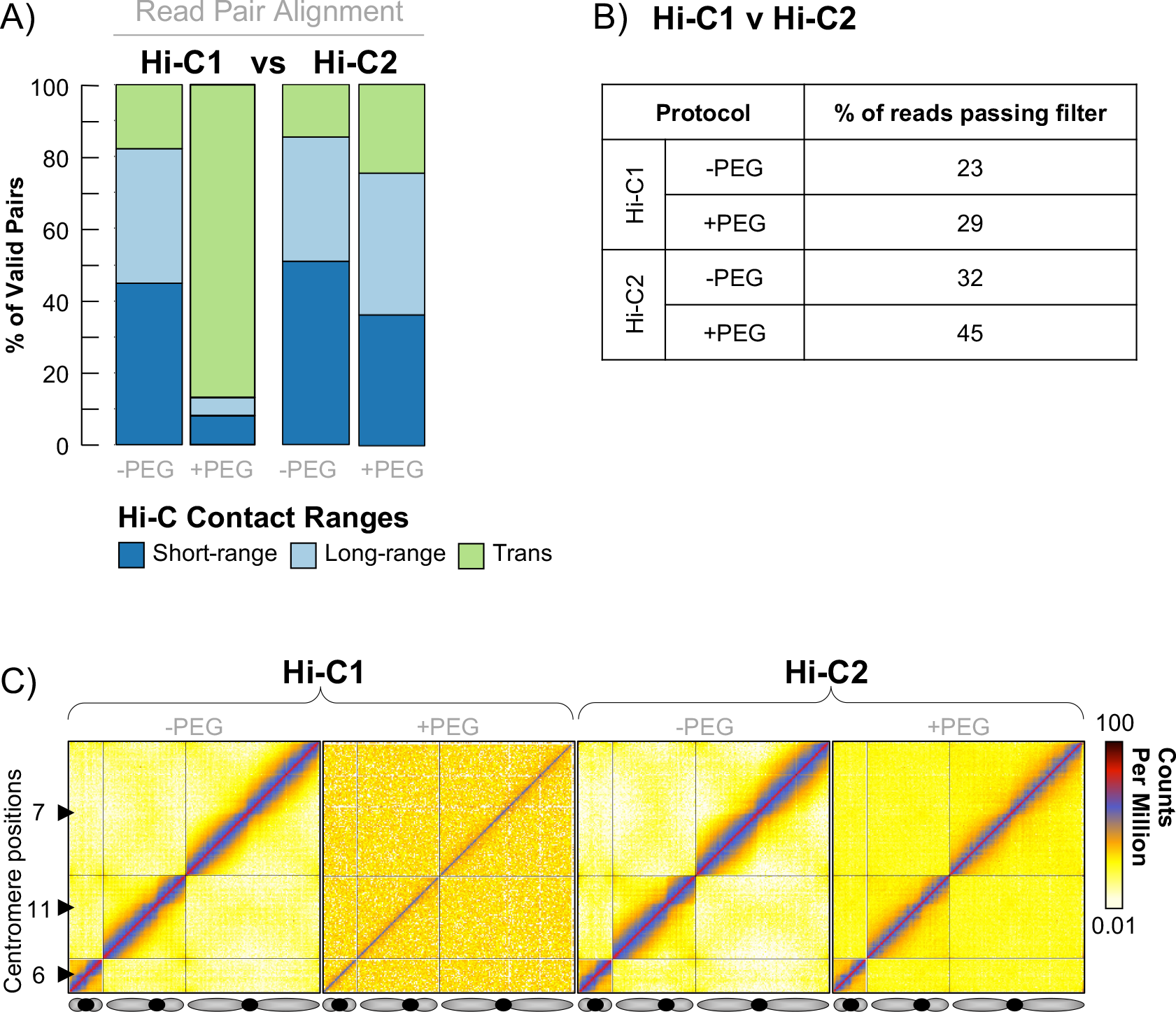
The effect of crowding agents during ligation stage. **A)** Fragment size distribution of short- and long-range intra-chromosomal (cis) contacts versus long-range inter-chromosomal (trans) contacts as reported by the Hi-C Pro pipeline *(14)* for libraries assayed by Hi-C1 ±PEG or Hi-C2 ±PEG (where Hi-C2 -PEG is equivalent to Hi-C2B, reported here). Addition of PEG-8000 to either protocol increases the rate of artifactual intermolecular ligation (detectable as increased trans contacts and higher matrix background (C), which is unwanted in Hi-C libraries). **B)** Table showing the percentage of reads that pass all Hi-C Pro filters as a fraction of the starting read value (all pairs) for Hi-C libraries assayed by Hi-C1 or Hi-C2 as described above. Increasing the rate of intermolecular ligation through the addition of PEG in either protocol generates an increase in the percentage of reads that pass all Hi-C Pro filters, but this increase in ligation efficiency is disproportionately made up of artifactual inter-molecular ligation products (detectable as increased trans contacts) at the expense of true intra-molecular proximity ligation events. **C)** Hi-C interaction maps from meiotic *S. cerevisiae* cells prepared by Hi-C1 or Hi-C2 as described above. Hi-C contact maps of chromosome 6, 11, and 7 are plotted at 10 Kb resolution. The black arrows indicate positions of the centromeres.
12. High levels of intermolecular ligation yield Hi-C libraries saturated with “noise”. To reduce the efficiency of the ligation reaction, the mixture is split across 16 PCR tubes, further reducing the relative frequency of non-specific intermolecular ligation.
13. Precipitated DNA should show evidence of white threads, or clouds; samples should be viscous, not solid.
14. A shorter spin can be used for the wash step. We use 5–10 minutes.
15. There should be two pellets per sample; the total volume will be ∼500 μL when pooled.
16. In our experience, between 1–5 μg of DNA per sample is sufficient to generate high resolution Hi-C libraries. Samples with <1 μg can be processed however; the library complexity is likely to be low. We recommend discarding samples with <1 μg.
17. For samples with >5 μg of DNA we suggest splitting the Hi-C sample between two separate reaction tubes for the dangling end removal step.
18. Hold any additional samples on ice during sonication.
19. To avoid cross contamination during NGS library preparation, filter tips should be used for all following steps.
20. The adapter ligation reaction can be run for more than 2 hours. We have performed the incubation for several hours or overnight without any effect on the final Hi-C library.
21. Between each wash aspirate the ethanol whilst keeping the tubes on the magnetic particle collector. Ensure the DNA/beads are completely dry before resuspending (eluting) in TLE following the second ethanol wash. Residual ethanol may interfere with downstream reactions.
22. In order to obtain high resolution Hi-C interaction maps, libraries should be sequenced to a depth of 10–30 million reads after passing all quality control filters. This usually equates to 50–100 million starting/unfiltered reads per Hi-C library. If wanting to quality control prior to deep sequencing, load small amounts of each library (aim for <1 million reads).
23. Default output generates ICED (balanced) and RAW matrices in addition to several other files including .bed files relating to each mapped resolution. Hi-C interaction data can be plotted using custom scripts (https://github.com/Neale-Lab/Hi-C-Scripts), or via standard packages (e.g., Hi-Glass: http://higlass.io/).

## Acknowledgements

We thank S. Schalbetter for her early assistance with this work, specifically, for implementing the 10-fold reduction of SDS concentration during the cell lysis step and production of “Hi-C1” libraries. This work is supported by Wellcome Trust Investigator (200843/Z/16/Z) and Discovery (225852/Z/22/Z) Awards to MJN.

